# Phenotypic distribution and selection response for linear-product gene-environment interaction in quantitative traits

**DOI:** 10.1101/2021.05.12.443944

**Authors:** Reginald D. Smith

**Affiliations:** Supreme Vinegar LLC, 3430 Progress Dr. Suite D, Bensalem, PA 19020

**Keywords:** GXE, gene-environment interaction, phenotypic plasticity, selection response

## Abstract

Gene-environment interaction is often described by linear phenotypic plasticity but has recently also been expressed as function of the product of genotype and environmental variables. While this model can be fitted in a multiple regression scenario, little has been written on the distribution of the product of breeding values and environment, *GE*, its expected moments, and the theoretical impact on phenotypic selection. Here we will explore these topics introducing the distribution for *GE*, its mean and variance, and its expected impact of lowering realized heritability due to is increasing the phenotypic variance.

## 1 Introduction

Gene-environment interaction has played an important, though not often well-known, role in the history of genetics (Hogben, 1933; Falconer, 1952; Robertson, 1959; Via and Lande, 1985; Mulder and Bijma, 2005; Mulder et. al., 2006). Gene-environment interaction is most often defined as a situation under which the contribution to phenotype by genotype varies across different environments. This posits a function *I*(*G*, *E*), where *G* is the genotype, most commonly represented as the breeding value, and *E* is the relevant environmental variable(s). This restates the traditional equation for contributions to phenotypic value as

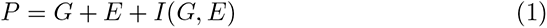

Gene-environment interaction is distinct from the normal additive effects of environment across all genotypes which varies phenotype with respect to a fixed mean breeding value, environment, as well as gene-environment covariance (GXE covariance). Unlike gene-environment interaction which is inherent in the organism under a given scope of genotypes and environments, GXE covariance can be eliminated by randomization of genotypes with respect to relevant environmental variables. A common model for gene-environment interaction is linear phenotypic plasticity (reaction norm) where the phenotypic average of a population varies linearly with the values of certain environmental variables the population is exposed to (Scheiner, 1993; Via et. al., 1995; De Jong, 1995; De Jong and Bijma, 2002; Kolmodin and Bijma, 2004; Mulder and Bijma, 2005).

There have been many studies of gene-environment interaction and its affect on selection with the linear phenotypic plasticity model (Van Tienderen and De Jong, 1994; Via et. al., 1995; De Jong, 1995; De Jong and Bijma, 2002; Kolmodin and Bijma, 2004; Mulder and Bijma, 2005; Mulder et. al., 2006), however, more recently new models of gene-environment interaction have found use, especially in medical genetics.

Guo (Guo, 2000) introduced a model where the gene-environment interaction is defined as the product of *G* and *E* along with an effect size constant. This work and later studies (Ma et. al., 2011) use multiple linear regression models of the form

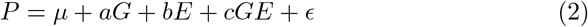

The gene-environment interaction is given by the *GE* term. Guo analyzed the properties of this gene-environment interaction for a single locus being affected by a single binary risk factor (environmental variable). For many quantitative traits with continuous values under the influence of many loci (aka the Fisher infinitesimal model (Barton et. al., 2017)), alternate techniques are necessary to describe the distribution of gene-environment interaction and its effect on the moments of the phenotype in the population as well as the response to selection.

## 2 Statistical distribution of gene-environment interaction in the linear-product model

The model we will define has phenotypic population mean *µ* and distributions of breeding values and environmental values defined as the probability of deviation from the mean value of the population. In other words, both *G* and *E* are normally distributed with means of zero and variances 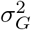 and 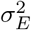. Also, the gene-environment interaction is represented by a term *c*_1_*GE* where *c*_1_ is an effect size constant. Thus the phenotype in the population is defined as

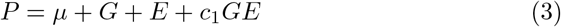

In order to determine the distribution and moments of the gene-environment interaction contribution, we rely on recent work defining the statistical distribution of the product of two normal variables. In recent years, an exact form for the distribution of the product of two normal variables has been derived by (Nadarajah and Pogány, 2016; Cui et. al., 2016) and also been related to the variance-gamma distribution (Gaunt, 2014, 2019, 2021). In particular, for two correlated normal distributions with mean 0, variances 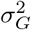 and 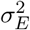, and gene-environment correlation *r*_*GXE*_ the probability density of their product *I* = *GE* is

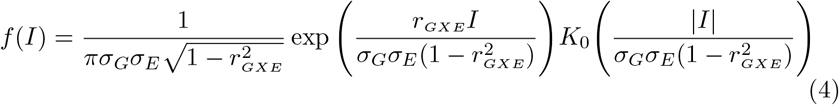

In the case where there is no GXE covariance and *r*_*GXE*_ = 0, this reduces to

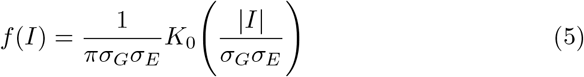

The term *K*_0_ in the two equations represents the modified Bessel function of the second kind of order zero. While not a widely familiar function, the moments for this distribution can be stated below by reference to the related variance-gamma distribution (Gaunt, 2014, 2019, 2021). Variance-gamma (*V G*) distributions are <u>in</u> the appendix, but the distribution 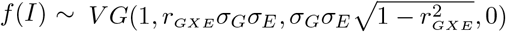 The first and second moments of this distribution are given by

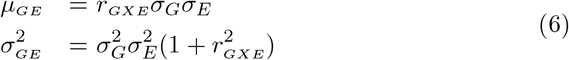

Again, when GXE covariance is absent

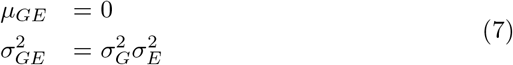

These moments lead to several conclusions about the effect of linear-product gene-environment interaction on the phenotypes of quantitative traits. In the absence of GXE covariance, there is no effect on the phenotypic mean. When covariance is present, however, the mean shifts proportionally to the value of the *GXE* covariance. In short the mean will shift by a value *c*_1_*σ*_*GXE*_.

The variance of *GE* will always widen proportionally to the product of the breeding value and environmental variances 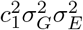, with the increase rising with gene-environment covariance to a maximum value proportional to twice the product of the breeding value variance, the environmental variance, and the effect size squared.

The overall phenotypic variance is directly affected only by the variance of *GE* since there is no covariance between either *G* nor *E* with *GE*. Covariances for product variables have been derived by (Bohrnstedt and Goldberger, 1969). From their equation 13, assuming *x* = *G, y* = *E, u* = 1, Δ *u* = 0, and *v* = *G*, we define *σ*_*GXGE*_ as

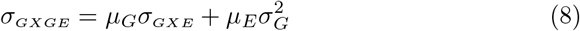

A similar derivation can be done for the covariance of *E* and *GE*. Given our parameterizations of *µ*_*G*_ = *µ*_*E*_ = 0, *σ*_*GXGE*_ = 0, thus the covariance of *G* or *E* with *GE* is zero.

In summary, product model *GXE* interaction will always widen the variance proportionally to the product of the variances of *G* and *E* but a population mean shift can only occur if there is simultaneous gene-environment covariance. This can be confirmed through numerical simulation where the first and second moments of populations with linear-product gene-environment interaction match theoretical expectations.

## 3 Phenotypic normality with the influence of *GE*

A key question is whether the product gene-environment interaction will induce a measurable distortion in the distribution of the phenotypic trait, which in most cases is distributed normally. The prior derivation of the first two moments is not conclusive to show this since higher order moments may deviate from normal expectations. If *G* or *E* and *GE* were statistically independent, the third moment for example would be given by 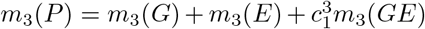.

The third and higher moments of *GE* are thus the deciding factors.

First, if *r* = 0, *GE* is symmetric and the third moment, and thus skew, are zero though excess kurtosis differs from normal distribution expectations. However, graphs of *GE* show that positive or negative gene-environment covariance induce right and left skews to the distributions respectively as shown in figures 2 and 3. Therefore, given a large enough value of *c*_1_, departures from normality can be expected under substantial gene-environment covariance.

**Figure 1:**
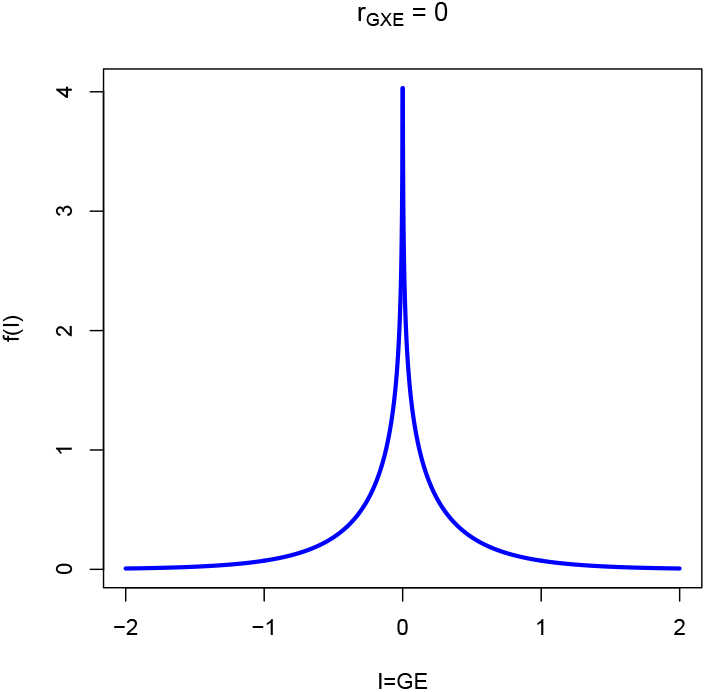
The probability density of *GE* for zero gene-environment correlation where 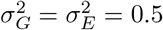.

**Figure 2:**
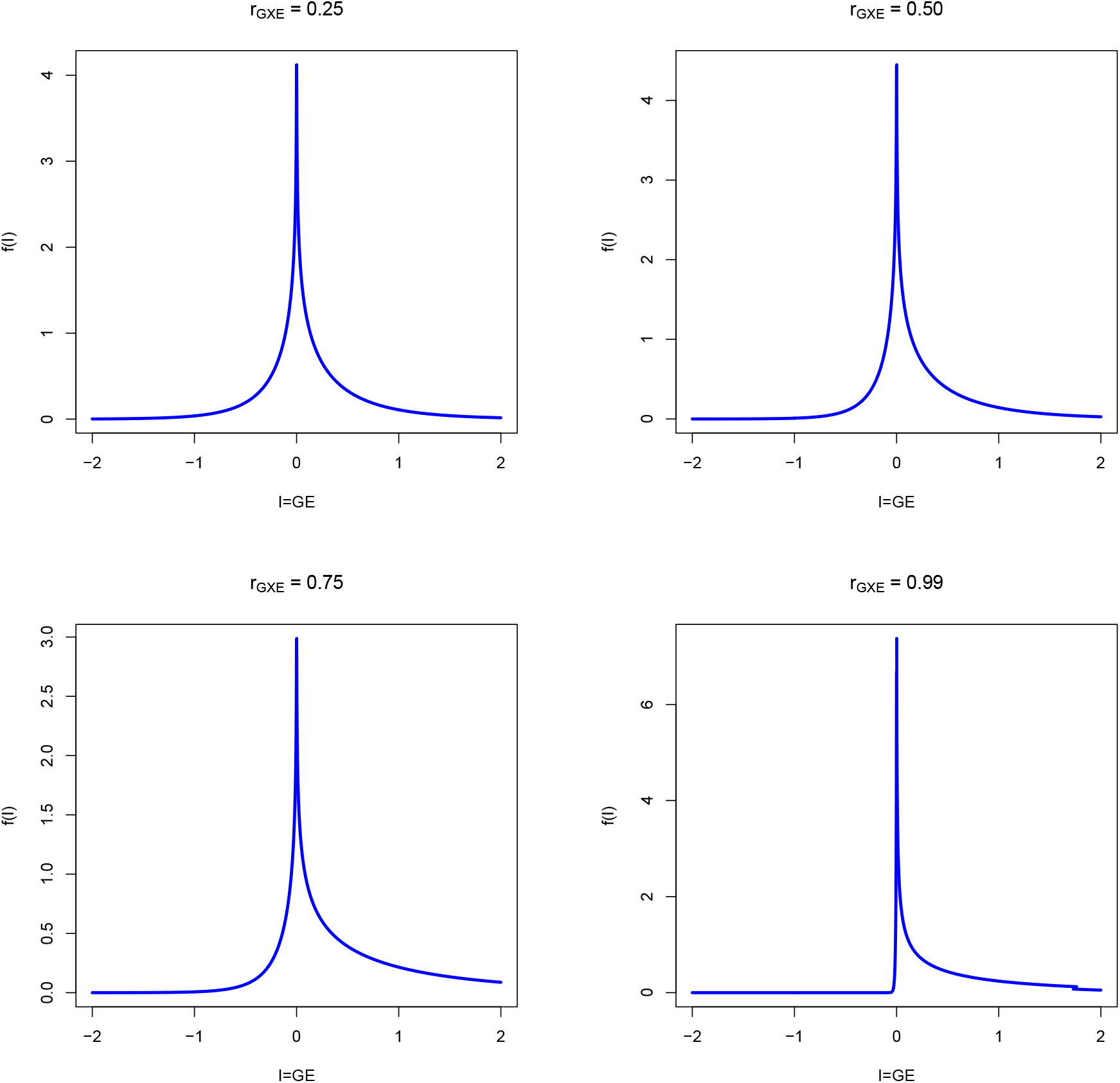
The probability densities of *GE* for various values of positive gene-environment correlation where 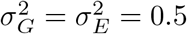.

**Figure 3:**
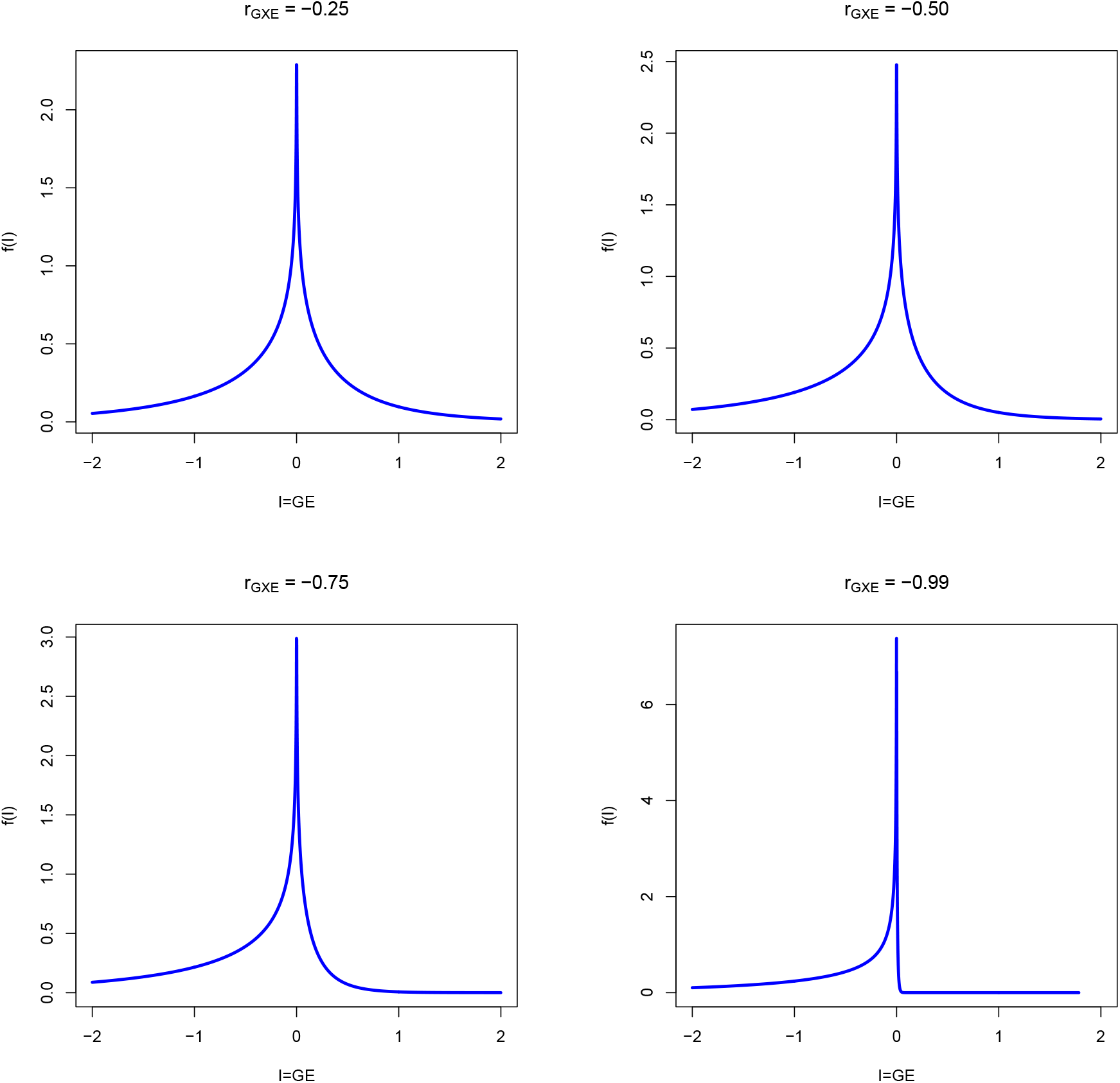
The probability densities of *GE* for various values of negative gene-environment correlation where 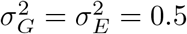.

However, even when *r* = 0, we will show the phenotypic distribution becomes skewed for large values of *c*_1_. While the covariance between *G* or *E* and *GE* indicates no linear dependence, the variables are not completely statistically independent from *GE*. There is nonlinear dependence which makes the simple addition of moments incorrect in calculating the higher order moments and produces skewed distributions.

In figure 4, for several values of *c*_1_ and *r* = 0 we show the phenotypic distribution and describe the calculated skew. Though the covariance is zero in all circumstances, the mutual information, which measures linear and nonlinear dependence is 0.4 between *G* and *GE*. As can be seen, when *c*_1_ becomes greater than approximately 0.25, substantial right skews develop in the phenotypic distribution due to the greater influence of the nonlinear dependence between the *G* or *E* and *GE*. Therefore, even if gene-environment covariance is considered absent, large effect sizes for *GE* would seem to dictate non-normality in the phenotypic distribution of the trait.

**Figure 4:**
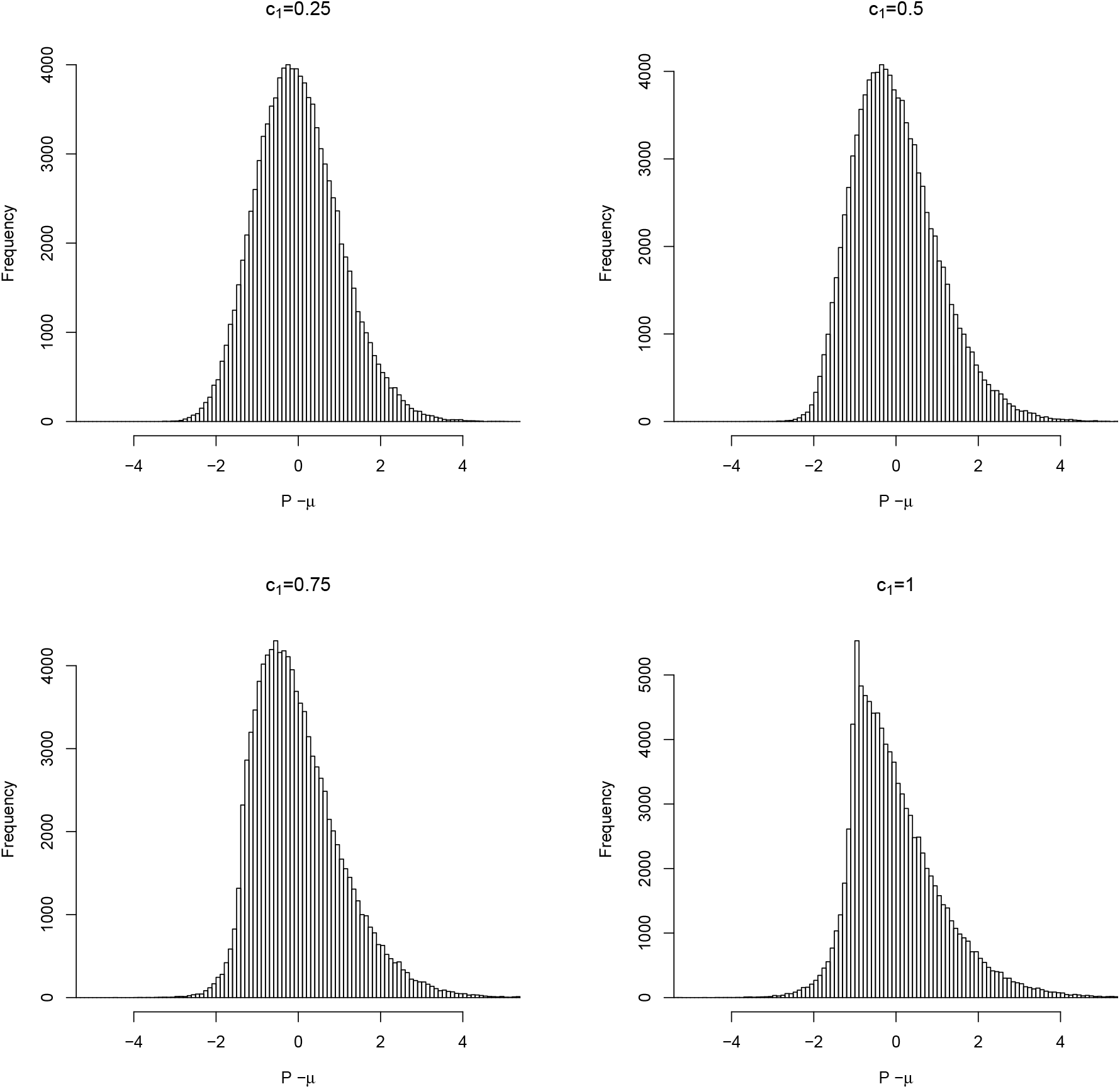
Graphs of the phenotypic distribution for a *N* = 100, 000 simulated distribution where gene-environment correlation *r*_*GXE*_ = 0 but where the effect size *c*_1_ of gene-environment interaction varies. Mutual information between *G* and *GE* is 0.4. Skew for each value of *c*_1_ are in parentheses: 0.25 (0.37), 0.5 (0.68), 0.75 (0.92), 1 (1.07).

## 4 Results

### 4.1 Linear-product gene-environment interaction and selection response

The previous discussion of phenotypic normality raises the question of how response to phenotypic selection plays out for linear-product model gene-environment interaction. Fortunately, as we will demonstrate, expectations for realized heritability are influenced little by the phenotypic skew and are largely in expectation with linear models.

We calculate the realized narrow heritability 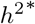 as defined in (Van Tienderen and De Jong, 1994) as

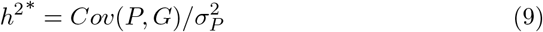

Though realized heritability is accurately defined using equation 9 only for those models with strictly linear dependence between phenotype and genotype, it can be used to investigate how the nonlinearity of this model affects a simulated selection response.

Given 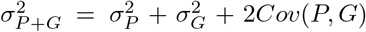 and 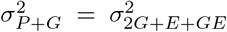 we can Calculate

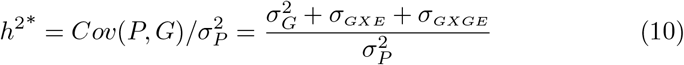

Therefore, the gene-environment interaction does not contribute to the covariance between phenotype and breeding value since the covariance between genotype and the genotype-environment product has already been shown to be zero. However, it does contribute to the realized heritability due to the phenotypic variance in the denominator and thus we can define

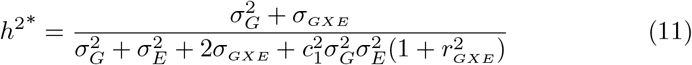

Thus, the only effect of product gene-environment interaction in the response of selection is to lower the realized narrow heritability due to the increase in phenotypic variance. Therefore, a slightly weaker selection response is expected. In Figure 5, a population of 100,000 individuals with randomly selected values of *G* and *E* and *GE* are subject to directional phenotypic selection at one standard deviation above the mean. The mean breeding values post-selection are divided by the difference between the phenotypic mean of the selected population and the mean of the original population(equation 6) to estimate realized heritability and compared against the calculation of equation 11. This is done for various combinations of *r*_*GXE*_ and *c*_1_ in Figure 5.

**Figure 5:**
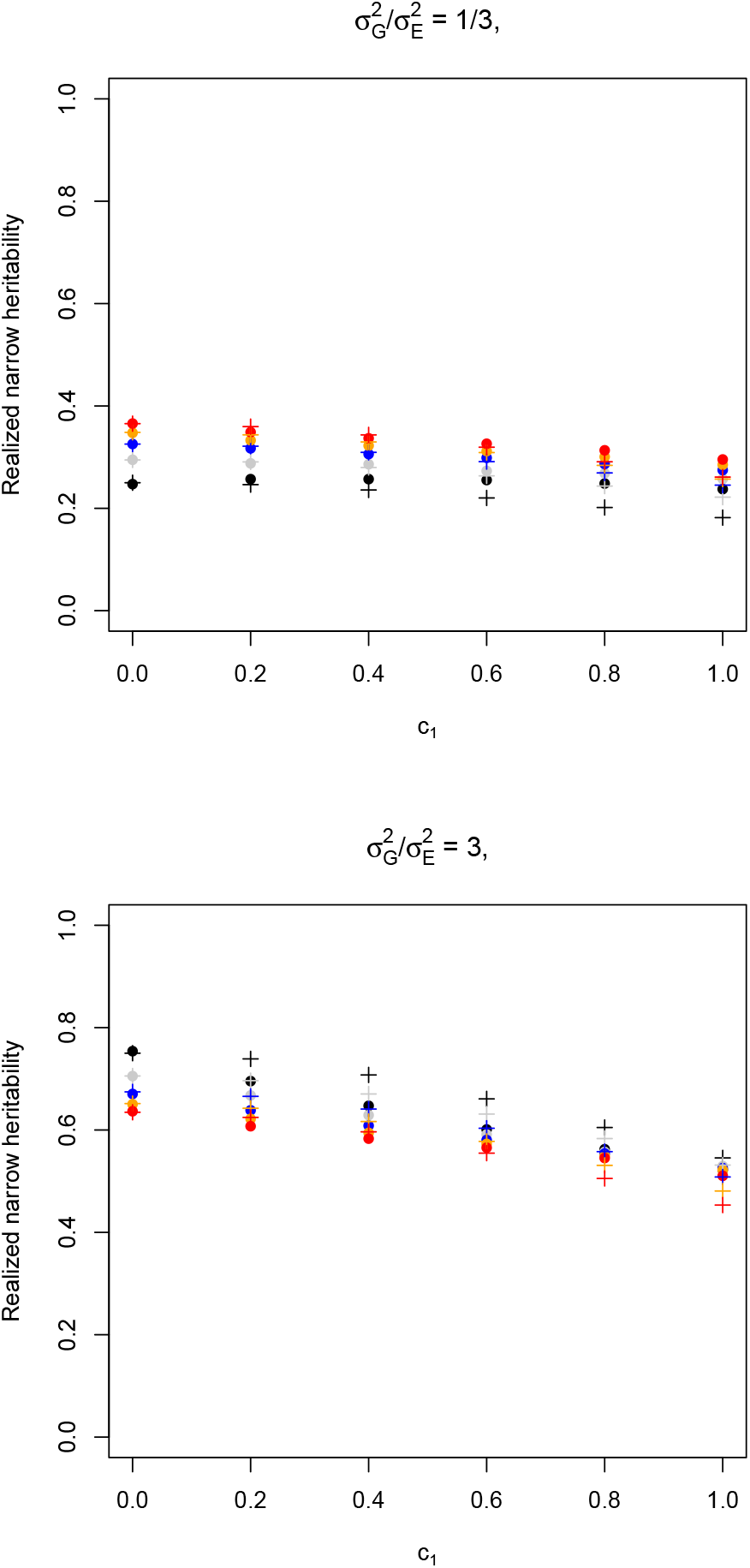
Realized values of 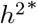 per theoretical values (crosses) and actual values (circles) of a *N* = 100, 000 point simulated population subject to positive directional phenotypic selection at *T* = *σ*, for various values of *r*_*GXE*_ and *c*_1_. Each color represents a different *r* where *r* ∈ (0, 0.25, 0.5, 0.75, 0.99) are black, grey, blue, orange, and red respectively.

The response to selection, measured through the realized heritability, does not deviate far from expected theoretical values. The largest deviation actually comes when there is no GXE covariance. Therefore, selection response models only accounting for the increase in phenotypic variance lowering the realized heritability will in most cases yield acceptably accurate results.

## 5 Discussion

Gene-environment interaction is an important aspect of explaining how the values and distributions of phenotypes emerge across a variety of populations and environments. While linear phenotypic plasticity is the most widely used and understood model for this interaction, interactions arising from the product of genetic and environmental factors have received increasing focus and interest in studies of gene-environment interaction. This paper has described the basic expected statistical aspects of such interactions and how they would change response under phenotypic selection. In summary, the variance of the population is increased proportional to the product of breeding value and environmental variances while the population mean is not changed except in the presence of GXE covariance. For normally distributed populations, the effect size measured by *c*_1_ is necessarily of low to moderate value if no unambiguous skew is evident in the distribution.

The effect of linear-product gene-environment interaction is clear in that by shifting the mean and widening the variance, a greater range of phenotypes should be present in the population than if interaction were absent. For threshold type traits which are often used to model diseases, etc. the proportion of the population above the threshold can raise significantly, especially if GXE covariance is also present.

The response to phenotypic selection is widely in line with what is expected due to the effects of a larger variance lowering realized heritability. The components that drive selection response, additive genetic variance and GXE covariance, see their impact unaffected by linear-product gene-environment interaction.

## 6 Data Availability Statement

No data was used for this paper.

## A Appendix

### The Variance Gamma Distribution

The variance gama distribution is a statistical distribution used in a variety of fields including economics and finance due to its importance in modeling stochastic phenomena, such as asset price returns, that exhibit skewed tails but finite moments in their statistical distributions. It has many forms but the one most useful in our analysis is due to Gaunt (2014, 2019, 2021). The variance gamma distribution is described by four parameters, *r, θ, σ*, and *µ* and has a formal expression of

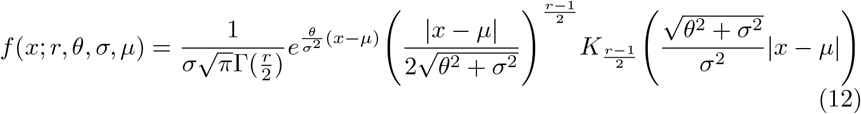

The distribution is often abbreviated *V G*(*r, θ, σ, µ*) and the parameters have values of *r >* 0, *θ* ∈ ℝ, *σ >* 0, and *µ* ∈ ℝ. Under certain parameterizations, the variance gamma distribution reduces to more common distributions. The distribution 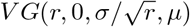 converges to the normal distribution *N* (*µ, σ*_2_) as *r* and *V G*(2, 0, *σ, µ*) is equivalent to the Laplace(*µ, σ*) distribution. The first two moments for the variance gamma distribution are given by Gaunt (2014)

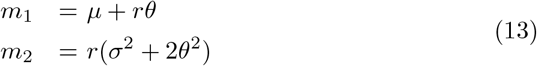

The distribution of the product of two normally distributed random variables both of mean zero, but with variances 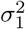 and 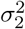 and which have a bivariate normal distribution of correlation, *<u>ρ</u>*<u>, ca</u>n be represented by the variance gamma distribution 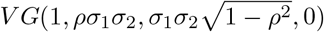.

## B References

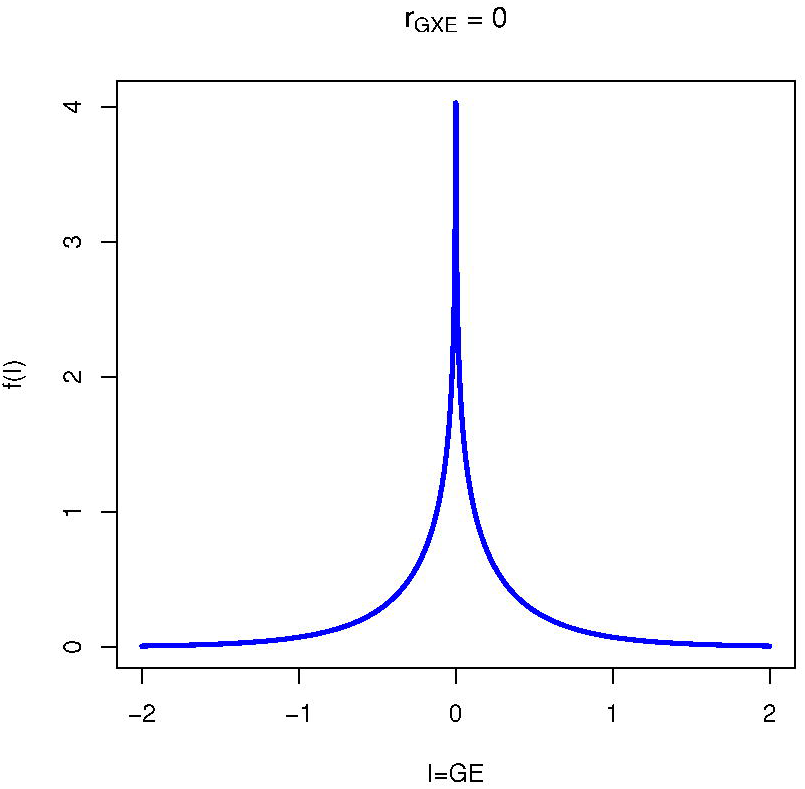

## References

Barton, NH, Etheridge, AM, Véber, A (2017) The infinitesimal model: Definition, derivation, and implications. Theor. Popul. Biol. 118: 50–73.

Bohrnstedt, GW, Goldberger, AS (1969). On the exact covariance of products of random variables. J. Am. Stat. Assoc. 64: 1439–1442.

Cui, G, Yu, X, Iommelli, S, Kong, L (2016) Exact distribution for the product of two correlated Gaussian random variables. IEEE Signal Process. Lett. 23: 1662–1666.

De Jong, G. (1995) Phenotypic plasticity as a product of selection in a variable environment. Am. Nat. 145: 493–512.

De Jong, G, Bijma, P (2002) Selection and phenotypic plasticity in evolutionary biology and animal breeding. Livest. Prod. Sci. 78: 195–214.

Falconer, DS (1952) The problem of environment and selection. Am. Nat. 86: 293–298.

Gaunt, RE (2014) Variance-Gamma approximation via Stein’s method. Electron. J. Probab. 19: 1–33.

Gaunt, RE (2019) A note on the distribution of the product of zero-mean correlated normal random variables. Stat. Neerl. 73: 176–179.

Gaunt, RE (2021) Stein’s method and the distribution of the product of zero mean correlated normal random variables. Commun. Stat. Theory Methods 50: 280–285.

Guo, SW (2000). Gene-environment interaction and the mapping of complex traits: some statistical models and their implications. Hum. Hered. 50: 286–303.

Hogben L (1933) Nature and Nurture (A reprint of William Withering Memorial Lectures on the methods of medical genetics). George Allen and Unwin Limited: London.

Kolmodin, R, Bijma, P (2004) Response to mass selection when the genotype by environment interaction is modelled as a linear reaction norm. Genet. Sel. Evol. 36: 1–20.

Ma, S, Yang, L, Romero, R, Cui, Y (2011). Varying coefficient model for gene–environment interaction: a non-linear look. Bioinformatics 27: 2119–2126.

Mulder, HA, Bijma P (2005). Effects of genotype x environment interaction on genetic gain in breeding programs. Anim. Sci. J. 83: 49–61.

Mulder, HA, Veerkamp, RF, Ducro, BJ, Van Arendonk, JAM, Bijma, P. (2006) Optimization of dairy cattle breeding programs for different environments with genotype by environment interaction. J. Dairy Sci. 89: 1740–1752.

Nadarajah, S, Pogány, TK (2016) On the distribution of the product of correlated normal random variables. Comptes Rendus Math. 354: 201–204.

Robertson, A (1959). The sampling variance of the genetic correlation coefficient. Biometrics 15: 469–485.

Scheiner, SM (1993) Genetics and evolution of phenotypic plasticity. Annu. Rev. Ecol. Evol. Syst. 24: 35–68.

Van Tienderen PH, De Jong G (1994) A general model of the relation between phenotypic selection and genetic response. J. Evol. Biol. 7: 1–12.

Via, S, Lande, R (1985) Genotype-environment interaction and the evolution of phenotypic plasticity. Evolution 39: 505–522.

Via, S, Gomulkiewicz, R, De Jong, G, Scheiner, SM, Schlichting, CD, Van Tienderen, PH (1995) Adaptive phenotypic plasticity: consensus and controversy. Trends Ecol. Evol. 10: 212–217.

